# Chromosome-scale genome assembly of B10.RIII, an autoimmune susceptible mouse strain

**DOI:** 10.1101/2025.05.16.654505

**Authors:** Vijayaraj Nagarajan, Yingyos Jittayasothorn, Guangpu Shi, Reiko Horai, Rachel R. Caspi

## Abstract

**Background:** A vast majority of potential drug candidates are initially tested in mice before human trials, resulting in life-saving treatments and preventive measures. Hundreds of autoimmune disorders, including arthritis, lupus etc., affect millions of people all over the world. Studying those disorders using mouse models is crucial for understanding the disease mechanisms. Knowing the genome sequence of these mouse models would help with those vital studies. B10.RIII is one of the most susceptible mouse strains used as a model to understand autoimmune diseases but did not have its genome sequenced until now.

**Findings:** In this study, we generated the first, high-quality chromosome level genome sequence of B10.RIII, using a combination of long read, short read and optical mapping techniques. The B10.RIII genome sequence scored higher than the reference mouse genome both in assembly completeness and assembly quality. The B10.RIII genome annotation based on the reference genome identified about 98% of the known reference genes.

**Conclusions:** We believe the availability of B10.RIII genome sequence will help advance the understanding of autoimmune diseases, giving the researchers a better idea on its unique genetic makeup. Comparative studies of B10.RIII with other mouse models could also significantly enhance the power of these models towards their clinical applications.

## Data Description

### Context

#### Importance of B10.RIII for the research community

Autoimmune diseases (ADs) are a diverse group of conditions resulting from immune disruptions causing aberrant B cell and T cell reactivity to normal human tissues. Over 80 ADs have been reported so far that affect approximately 5%–8% of the world population, causing a huge global socioeconomic issue [1]. ADs can be systemic, such as rheumatoid arthritis (RA), systemic lupus erythematosus (SLE), and ankylosing spondylitis (AS) damaging multiple organs; or they can be organ-specific, being restricted to a specific organ or tissue, such as multiple sclerosis, autoimmune uveitis, and type 1 diabetes affecting the central nervous system, eye, and pancreas, respectively [2, 3]. Although studies in the past decades have provided increasingly sophisticated knowledge about human ADs, the detailed mechanisms remain poorly understood [4]. For treatment, the immunomodulatory drugs currently used for ADs are not disease-specific and can cause severe side effects such as infections and malignancies [2, 3]. Animal-based lab research is urgently needed to explore the detailed mechanisms leading to human ADs in pursuance of effective treatment and even prevention. Among lab animals, mice are the most widely used organisms because of their genetic and physiological homologies to humans. It’s estimated that about 99% of mouse genes have a homolog in the human genome, and about 80% of the human and mouse genomes reside within conserved syntenic segments [5].

The classic method to reproduce human ADs in the mouse is to immunize the animal using tissue-specific peptides or proteins in combination with adjuvants [6]. This has been a standard protocol for many experimental mouse models, such as experimental autoimmune encephalomyelitis (EAE), uveitis (EAU), neuritis (EAN), thyroiditis (EAT), and experimental collagen-induced arthritis (ECIA) [7–12]. Significant differences among mouse strains in their susceptibility to disease induction have been well documented, with B10.RIII being the most susceptible one in many cases [8, 13–15]. These variations are found depending on mouse background and MHC haplotype, both of which are indispensable. For example, the permissive backgrounds for EAU induction are B10.RIII, B10.BR, B10.A, C57BL/6, and C57BL/10, while the susceptible haplotypes are H-2r, H-2k, and H-2b [16]. Genetic interactions play an important role in driving these autoimmune processes. Given its high susceptibility and unique disease time-course, the B10.RIII mouse has served as an invaluable tool in lab studies; however, the whole genome assembly of B10.RIII is currently lacking. The inbred mouse strain C57BL/6 (B6) is the most cited and well-characterized laboratory strain used in biomedical research [17]. Its *de novo* genome assembly was finished in 2009 [18]. Several groups have been working on whole genome sequencing (WGS) for other strains. In 2018, full-length draft *de novo* genome assemblies for 16 widely used inbred mouse strains have been completed [19].

In this article, we report a high-quality chromosome-scale genome assembly of the B10.RIII mouse strain built using three different sequencing platforms, including Illumina sequencing, Single Molecular, Real-Time (SMRT) sequencing (PacBio), and Bionano Optic Mapping. The genomic information reported here will be greatly beneficial for accelerating animal studies in the field of AID.

## Methods

### Mouse DNA Extraction

The female B10.RIII (B10.RIII-*H2^r^ H2-T18^b^*/(71NS)SnJ) mouse was purchased from The Jackson Laboratory (ME, USA, strain #: 000457). The animal was maintained under specific-pathogen free (SPF) conditions and euthanized at 13 weeks of age by Carbon dioxide (CO2) inhalation. Splenocytes were isolated using 1X PBS (Gibco), ACK lysing Buffer (Quality Biological), FCS serum (HyClone), RPMI medium (HyClone), DNA Stabilizer (Bionano Genomics), and Cell Buffer (Bionano Genomics). Single cell suspension of 6.8 million Splenocytes were used for high-molecular-weight genomic DNA (HMW gDNA) extraction, using MagAttract HMW DNA Kit from Qiagen (Cat. No. 67563) for PacBio. Ultra-high molecular weight (UHMW) gDNA was extracted from mouse splenocytes with the Bionano Prep SP Blood and Cell DNA Isolation Kit (Bionano Genomics, 80030) as described in the Bionano Prep SP Frozen Cell Pellet DNA Isolation Protocol (Bionano Genomics). Briefly, 5 million frozen pelleted cells were thawed in a 37°C bath and resuspended in DNA Stabilizing Buffer (Bionano Genomics). After that, the cells were lysed in the presence of detergents, proteinase K, and RNase A and the UHMW gDNA was bound to a silica Nanobind Disk (Bionano Genomics), washed, and eluted. The extracted gDNA was equilibrated for 3 days at room temperature to homogenize and quantified with Qubit dsDNA BR assay kit (Thermo Fisher Scientific, Q32850) and Femto Pulse (Agilent). The HMW gDNA was used for both the PacBio and Illumina Sequencing.

### Illumina sequencing

Illumina TruSeq Nano DNA Library Prep kit was used to prepare uniquely indexed paired-end (2x150bp) libraries of genomic DNA (gDNA). Briefly, the genomic DNA was fragmented to ∼ 500 bp insert size on the Covaris which generates dsDNA fragments with 3’ or 5’ overhangs. The sheared DNA is blunt-ended, and library size selection done using sample purification beads. A single ‘A’ nucleotide is added to the 3’ ends of the blunt fragments to prevent them from ligating to each other during the adapter ligation reaction. A corresponding single ‘T’ nucleotide on the 3’ end of the adapter provides a complementary overhang for ligating the adapter to the fragment. The indexed adapters are ligated to the ends of the DNA fragments and then PCR-amplified to enrich for fragments that have adapters on both ends. The final purified product is then quantitated by qPCR before cluster generation and sequenced using NovaSeq sequencing platform.

### PacBio sequencing

The PacBio library was prepared following the “HiFi SMRTbell® Libraries using the SMRTbell Express Template Prep Kit 2.0” protocol (Pacific Biosciences, CA). Briefly, gDNA was sheared to ∼17 kb using the Megaruptor 3 instrument (Diagenode, Inc.). Single-stranded overhangs removal, DNA damage repair, end-repair/A-tailing, adapter ligation, nuclease treatment and AMPure PB bead purification steps were performed according to manufacturer’s recommendations. A size selection was performed using the BluePippin system (Sage Science) to remove library fragments shorter than 10 kb. Sequencing primer v5 was annealed and Sequel II polymerase 2.2 was bound to the library prior to loading on four SMRTcells 8M on the PacBio Sequel II System using adaptive loading. Sequencing was performed for 30 h.

### Bionano sequencing

The isolated UHMW gDNA was fluorescently labeled using the Bionano Prep DNA Labeling Kit-DLS (Bionano Genomics, 80005) according to the Prep Direct Label and Stain (DLS) Protocol (Bionano Genomics). In short, Direct Label Enzyme (DLE-1) and DL-green fluorophores were used to label 750 ng of purified gDNA at a specific sequence motif. After a cleanup of the fluorophores excess, DNA backbone was counterstained overnight before quantitation with Qubit dsDNA HS assay kit (Thermo Fisher Scientific, Q32851). Finally, the labeled UHMW gDNA molecules were loaded on a Saphyr G2.3 chip for sequentially imaging across nanochannels on the Saphyr instrument (Bionano Genomics).

### K-mer estimation

KmerGenie [20] was used to estimate the best k-mer length, with the command; “`kmergenie inputs -t60`”. The ‘inputs’ variable stored the list of raw Illumina fastq files.

### Genome size estimation

Genome size was estimated using the k-mer counting tool Jellyfish [21], with the commands; “`jellyfish count -t 60 -C -m 117 -s 30G -o jelly.jf -F 2 <(zcat F1_S1_R1_001.fastq.gz) <(zcat F1_S1_R2_001.fastq.gz)`”. The ‘k’ of 117 was used with Jellyfish, based on the previous KmerGenie prediction of the best k. Jellyfish stats and histo commands were used to identify the total amount of k-mers and peak k-mer size.

Based on the k-mer histogram data, the formula of “`sum(as.numeric(jel[5:9989,1]*jel[5:9989,2]))/18`” was used to estimate the genome size. Jellyfish data was analyzed in Rstudio, using the included jellyanalysis.R script.

### Genome properties estimation

GenomeScope [22] was used with the Jellyfish k-mer frequency count as the input, to estimate genome characteristics based on illumina short reads, with k-mer_length of 21 and read_length of 150.

### Short read preprocessing

Quality of the Illumina short reads were accessed using the fastqc [23] tool. BBDUK from the BBTools suite [24] was used to check for adapter contamination with the command “`bbtools bbduk in1=F1_S1_R1_001.fastq.gz in2=F1_S1_R2_001.fastq.gz k=23 ref=adapters.fasta stats=bbdukstats.txt”`. Adapters were removed using the command “bbtools bbduk in1=F1_S1_R1_001.fastq.gz in2=F1_S1_R2_001.fastq.gz out1=F1_S1_R1_001.trimmed.fastq.gz out2=F1_S1_R2_001.trimmed.fastq.gz ktrim=r k=21 mink=11 hdist=1 ref=adapters.fasta stats=bbduk-trim-stats.txt 2> log.txt”. Trimmed adapter removed fastq files were again checked with bbduk to make sure no contamination existed.

### Long read preprocessing

PacBio HiFi reads were preprocessed to remove adapter contamination using the HiFiAdapterFilt [25] tool, with default parameters.

### Genome assembly

Primary assembly of the PacBio HiFi data was carried out using both raw and filtered reads. HiCanu [26], IPA [27] and Hifiasm [28] were used for the primary assembly. HiCanu was run with the command “`canu -p hicanudefault3 -d hicanudefault3 genomeSize=2.7g -pacbio-hifi *.fastq.gz batThreads=24 batMemory=247g`”. IPA was run with the command “`ipa local --nthreads 20 -- njobs 1 --run-dir defaultfiltered -i F1_1.ccs.filt.fastq.gz -i F1_2.ccs.filt.fastq.gz -I F1_3.ccs.filt.fastq.gz -i F1_4.ccs.filt.fastq.gz`”. Hifiasm was run with the command “`hifiasm -o default-60-filtered.asm -t 200 F1_1.ccs.filt.fastq.gz F1_2.ccs.filt.fastq.gz F1_3.ccs.filt.fastq.gz F1_4.ccs.filt.fastq.g`z”.

### Assembly alignment for polishing

HiFi reads were aligned against the assembled genome using pbmm2 [29], with the command; “pbmm2 align asmdefaultfiltered.bp.p_ctg.fa subreadbams.fofn asmfilteredaligned-norm.bam --sort -j 56 -J 30 -m 8G --preset SUBREAD --log-level INFO --log-file asmdefaultfiltered-norm.log”

### Contig specific alignment for polishing

SAMtools faidx [30] was used to generate a file containing the list of contig names from the primary assembly. Alignment for each of the contigs were extracted from the full genome aligned bam file using bamtools merge [31] with region option. An example command used for generating contig specific alignment file for contig ptg000001l is “`bamtools merge -in asmfilteredaligned-norm.bam -out ptg000001l.bam -region ptg000001l`”

### Polishing with Arrow

gcpp [32] windows option was used to polish each of the 129 contigs in parallel. An example command used for polishing contig ptg000001l is “`gcpp -w ptg000001l:0-72362188 -r asmdefaultfiltered.bp.p_ctg.fa -o ptg000001l.polished.fasta ptg000001l.bam --log-level TRACE --log-file ptg000001l.log`”.

Polished contigs were merged to generate the polished assembly, using the command `cat`

*polished.fasta > arrow-polished-1.fasta.

### Assembly polishing using illumina reads

Pilon [33] was used for the second round of polishing with Illumina whole genome short reads. First the arrow polished assembly was indexed using the command `bwa index arrow- polished-1.fasta`. Trimmed and adapter filtered Illumina short reads were aligned using the following command: `bwa mem -t 120 arrow-polished-1.fasta F1_S1_R1_001.trimmed.fastq.gz F1_S1_R2_001.trimmed.fastq.gz -o bwa_mapping_illumina_for_pilon_on_arrow_polished-1.sam`. The aligned file was split for each of the contigs, sorted and indexed for parallel computation. Polishing was done for each of the contigs and then polished contigs were combined. An example command used for splitting the contig, sorting, indexing and polishing is: bamtools merge -in bwa_mapping_illumina_for_pilon_on_arrow_polished-1_sorted.bam - out ptg000001l\|arrow.bam -region ptg000001l\|arrow ; samtools sort -o ptg000001l\|arrow_sorted.bam ptg000001l\|arrow.bam ; samtools index ptg000001l\|arrow_sorted.bam ; java -Xmx1543168m -jar pilon-1.24.jar --genome ptg000001l\|arrow.fa --bam ptg000001l\|arrow_sorted.bam --output ptg000001l\|arrow --outdir testpilon. The polished individual contigs were combined using the command `cat *arrow.fasta > ../../pilon-parallel/arrow-1-pilon-1-polished- 1.fasta`.

### Hybrid scaffolding with Bionano data and polished contigs

Bionano’s HybridScaffold [34] was used to scaffold the assembly polished with arrow alone and also the assembly polished with arrow and then with pilon, using the following example command; perl hybridScaffold.pl -n arrow-1-pilon-1-polished-

1.fasta -b EXP_REFINEFINAL1.cmap -c hybridScaffold_DLE1_config.xml -r RefAligner -o arrow-pilon- polished -f -g -B 2 -N 2.

### Chromosome scaffolding

We performed GRCm39 (https://www.ncbi.nlm.nih.gov/datasets/genome/GCF_000001635.27/) reference-based chromosome scaffolding using chromosome_scaffolder.sh from MaSuRCA-4.1.0 package [35] and RagTag 2.1.0 [36], on three different versions of the assembly (unpolished-scaffold.fasta, arrow-polished-scaffold.fasta, arrow-pilon-polished-scaffold.fasta). An example command for MaSuRCA run is `bash chromosome_scaffolder.sh -r`

GCF_000001635.27_GRCm39_genomic.fna -q unpolished-scaffold.fasta

`-t 120 -nb`. An example command for RagTag run is ragtag.py scaffold GCF_000001635.27_GRCm39_genomic.fna unpolished-scaffold.fasta -t 120 -o unpolished.

### BUSCO analysis

The quality of the chromosome builds was analyzed using BUSCO suite, using glires_odb10 and the --auto-lineage-euk options for lineage reference. An example command is “`busco -c`

120 -m genome -l glires_odb10 -i GCF_000001635.27_GRCm39_genomic.fna.arrow-polished- scaffold.fasta.split.reconciled.fa -o arrow-masurca-gliers-mm39”.

### Annotation

A reference-based preliminary annotation was performed with the liftoff [37] tool using the command liftoff -g GCF_000001635.27_GRCm39_genomic.gff -o b10.gff3 -polish -chroms chromosomes.txt -f feature_types.txt - copies -p 55 -m minimap2 GCA_030265425.1_NEI_Mmus_1.0_genomic.fna GCF_000001635.27_GRCm39_genomic.fna. Annotation statistics were generated using AGAT suite [38], with the command agat_sp_statistics.pl --gff b10_swarm_polished.gff3.

### Data validation and quality control

Quality of the primary assemblies generated by three different assemblers, for both raw and filtered HiFi reads, were assessed using QUAST [39], along with the MM10 reference genome (GCF_000001635.27_GRCm39_genomic.fna), using the command `quast.py -o hicanu- pbipa-hifiasm hicanudefault4.contigs.fasta final.p_ctg.fasta default.asm.bp.p_ctg.fa GCF_000001635.27_GRCm39_genomic.fna -- threads 120.` The quality of the assemblies was also analyzed using QUAST after each of the steps, including polishing with long reads, polishing with short reads, hybrid scaffolding with Bionano data and chromosome scaffolding.

### Raw data summary

The raw data summary, including the total number of sequenced reads before and after quality control, for the long and short read sequencing are provided in Table 1, along with the bionano optical mapping summary.

**Table 1.** Raw data summary for PacBio, Illumina and Bionano data.

| PacBio long read data summary |  |
| --- | --- |
| Total number of bases sequenced | 2,790,916,468 |
| Total number of raw reads | 7,434,073 |
| Total number of reads after adapter removal | 7,432,281 |
| % GC | 41.43 |
| Illumina short read data summary |  |
| Total number of raw reads | 1,685,341,136 |
| Total number of reads after adapter removal | 1,684,745,048 |
| Read length | 151 bp |
| % GC | 41 |
| % Duplicate reads | 21.2 |
| Bionano optical map raw data summary |  |
| Total number of molecules | 23,131,891 |
| Total length | 2,742,065.31 Mbp |
| Average length | 118.54 Kbp |
| Molecule N50 | 230.25 Kbp |
| Label density (/100kb) | 18.93 |
| Median length | 63,000 |

### Genome size estimation

KmerGenie estimated the best k as 117. A total 48.8G of k-mers and the peak at 18 were identified using Jellyfish, providing an estimated genome size of 2.7 Gb. GenomeScope estimated a maximum heterozygosity of 0.157%, which is expected due to B10.RIII being extensively inbred. The KmerGenie report, Jellyfish histogram and the GenomeScope summary file are provided as supplementary files.

### Primary genome assembly

We performed the primary genome assembly of the PacBio HiFi reads, using three different assemblers. Assembly was carried out both before and after filtering the HiFi reads for adapters. We assessed the quality of the assembled genomes and identified that the Hifiasm_filtered primary assembly had the highest N50 score of 48307382, highlighted in bold in Table 2. The Hifiasm_filtered primary assembly was used for the further analysis.

**Table 2.** Primary genome assembly quality.

|  | HiCanu | HiCanu_filtered | IPA | IPA_filtered | Hifiasm | Hifiasm_filtered | GRCm39 |
| --- | --- | --- | --- | --- | --- | --- | --- |
| # contigs (>= 0 bp) | 6420 | 5196 | 2655 | 2656 | 116 | 129 | 61 |
| # contigs (>= 1000 bp) | 6420 | 5196 | 2655 | 2656 | 116 | 129 | 61 |
| # contigs (>= 50000 bp) | 513 | 518 | 1233 | 1233 | 98 | 101 | 38 |
| Total length (>= 0 bp) | 3069667042 | 3036410063 | 2753118178 | 2752968625 | 2756964303 | 2757347189 | 2728222451 |
| Total length (>= 1000 bp) | 3069667042 | 3036410063 | 2753118178 | 2752968625 | 2756964303 | 2757347189 | 2728222451 |
| Total length (>= 50000 bp) | 2899252771 | 2898976624 | 2708685009 | 2708491461 | 2756414576 | 2756602245 | 2727603288 |
| # contigs | 6420 | 5196 | 2655 | 2656 | 116 | 129 | 61 |
| Largest contig | 86262123 | 84883328 | 78529241 | 78528261 | 163900417 | 163900415 | 195154279 |
| Total length | 3069667042 | 3036410063 | 2753118178 | 2752968625 | 2756964303 | 2757347189 | 2728222451 |
| GC (%) | 41.22 | 41.21 | 41.32 | 41.32 | 41.43 | 41.43 | 41.67 |
| N50 | 34671040 | 32898314 | 26427946 | 26428279 | 48307378 | **48307382** | 130530862 |
| N90 | 2610807 | 3395394 | 4099099 | 4099105 | 20910107 | 19548717 | 95294699 |
| auN | 38004717.8 | 34973578.4 | 31344433.8 | 31344368.1 | 59062679.2 | 58979945.2 | 137559569.6 |
| L50 | 28 | 30 | 31 | 31 | 18 | 18 | 9 |
| L90 | 127 | 128 | 116 | 116 | 50 | 50 | 18 |
| # N's per 100 kbp | 0 | 0 | 0 | 0 | 0 | 0 | 2697.75 |

Quast genome assembly quality assessment of different primary assemblies, using both raw and filtered HiFi reads shown in the above Table 2. All statistics are based on contigs of size >= 500 bp, unless otherwise noted.

### Assembly polishing

The Hifiasm_filtered primary assembly was polished with raw HiFi reads, followed by polishing with adapter removed short reads. The long read based assembly polishing improved the N50 of the primary assembly, highlighted in Table 3. The long read based polished assembly was used for further analysis.

**Table 3.** Polished genome assembly quality.

|  | primary-assembly-unpolished | arrow-polished | arrow-pilon-polished | GRCm39 |
| --- | --- | --- | --- | --- |
| # contigs (>= 0 bp) | 129 | 129 | 129 | 61 |
| # contigs (>= 1000 bp) | 129 | 129 | 129 | 61 |
| # contigs (>= 50000 bp) | 101 | 102 | 102 | 38 |
| Total length (>= 0 bp) | 2757347189 | 2757544570 | 2757431338 | 2728222451 |
| Total length (>= 1000 bp) | 2757347189 | 2757544570 | 2757431338 | 2728222451 |
| Total length (>= 50000 bp) | 2756602245 | 2756847535 | 2756735904 | 2727603288 |
| # contigs | 129 | 129 | 129 | 61 |
| Largest contig | 163900415 | 163911670 | 163928538 | 195154279 |
| Total length | 2757347189 | 2757544570 | 2757431338 | 2728222451 |
| GC (%) | 41.43 | 41.43 | 41.43 | 41.67 |
| N50 | 48307382 | **48311567** | 48309103 | 130530862 |
| N90 | 19548717 | 19550139 | 19549361 | 95294699 |
| auN | 58979945.2 | 58984337.7 | 58984004 | 137559570 |
| L50 | 18 | 18 | 18 | 9 |
| L90 | 50 | 50 | 50 | 18 |
| # N's per 100 kbp | 0 | 0 | 0 | 2697.75 |

Quast genome assembly quality assessment of different primary assemblies after polishing with long reads alone (Arrow) and after polished with both long reads (Arrow) and short reads (Pilon), shown in the above Table 3.

### Assembly scaffolding

The Hifiasm filtered, arrow polished assembly was scaffolded with Bionano optical data, to build a hybrid assembly. The improved N50 of the polished and hybrid scaffolded assembly is highlighted in bold in Table 4.

**Table 4.** Polished hybrid scaffolded genome assembly quality.

|  | unpolished-scaffolded | arrow-polished_scaffolded | GRCm39 |
| --- | --- | --- | --- |
| # contigs (>= 0 bp) | 53 | 51 | 61 |
| # contigs (>= 1000 bp) | 53 | 51 | 61 |
| # contigs (>= 50000 bp) | 53 | 51 | 38 |
| Total length ($\geq 0$ bp) | 2790533087 | 2790915468 | 2728222451 |
| Total length ($\geq 1000$ bp) | 2790533087 | 2790915468 | 2728222451 |
| Total length ($\geq 50000$ bp) | 2790533087 | 2790915468 | 2727603288 |
| # contigs | 53 | 51 | 61 |
| Largest contig | 196192695 | 196210478 | 195154279 |
| Total length | 2790533087 | 2790915468 | 2728222451 |
| GC (%) | 41.45 | 41.45 | 41.67 |
| N50 | 137570235 | **137580593** | 130530862 |
| N90 | 71062120 | 71067871 | 95294699 |
| auN | 127959678.5 | 127963698 | 137559570 |
| L50 | 9 | 9 | 9 |
| L90 | 20 | 20 | 18 |
| # N's per 100 kbp | 1753.32 | 1753.27 | 2697.75 |

Quast genome assembly quality assessment of different primary assemblies after polishing with long reads alone (Arrow) and hybrid scaffolding with the Bionano optical map, shown in the above Table 4.

### Chromosome scaffolding

The hybrid assembly was chromosome scaffolded using two different tools, RagTag and MaSuRCA. The scaffolded chromosomes were assessed for their quality using Quast. The Arrow polished and RagTag scaffolded chromosome assembly had a high N50, as highlighted in bold in Table 5.

**Table 5.** Quality of the chromosome scaffolded genome assembly.

|  | arrow-polished-masurca | arrow-polished-ragtag | GRCm39 |
| --- | --- | --- | --- |
| # contigs ($\geq 0$ bp) | 41 | 41 | 61 |
| # contigs ($\geq 1000$ bp) | 41 | 41 | 61 |
| # contigs ($\geq 50000$ bp) | 41 | 41 | 38 |
| Total length ($\geq 0$ bp) | 2697335745 | 2790916468 | 2728222451 |
| Total length ($\geq 1000$ bp) | 2697335745 | 2790916468 | 2728222451 |
| Total length ($\geq 50000$ bp) | 2697335745 | 2790916468 | 2727603288 |
| # contigs | 41 | 41 | 61 |
| Largest contig | 195990569 | 196210478 | 195154279 |
| Total length | 2697335745 | 2790916468 | 2728222451 |
| GC (%) | 41.48 | 41.45 | 41.67 |
| N50 | 137607371 | **144428894** | 130530862 |
| N90 | 96149225 | 96154628 | 95294699 |
| auN | 134306017.5 | 140475790.7 | 137559569.6 |
| L50 | 9 | 9 | 9 |
| L90 | 18 | 18 | 18 |
| # N's per 100 kbp | 4.49 | 1753.31 | 2697.75 |

Quast quality assessment of chromosome scaffolded genome assemblies, using two different tools, shown in the above Table 4.

### BUSCO analysis

The BUSCO analysis for the completeness of the assembly shows our assembly of B10.RIII with less gaps, less missing BUSCOs and more complete BUSCOs, as shown in Table 5. The final assembly statistics are shown in Table 6.

**Table 5.** Busco analysis for genome completeness.

|  | **B10.RIII** | C57BL/6J (mm39, GRCm39) |
| --- | --- | --- |
| Complete BUSCOs (C) | 13306 | 13304 |
| Complete and single-copy BUSCOs (S) | 12912 | 12910 |
| Complete and duplicated BUSCOs (D) | 394 | 394 |
| Fragmented BUSCOs (F) | 86 | 85 |
| Missing BUSCOs (M) | 406 | 409 |
| Total BUSCO groups searched | 13798 | 13798 |
| Percent gaps | 1.75% | 2.70% |

**Table 6.** The final assembly statistics for B10.RIII, along with the reference genome assembly statistics.

|  | B10.RIII | C57BL/6J (mm39, GRCm39) |
| --- | --- | --- |
| Genome size | 2.8 Gb | 2.7 Gb |
| Total ungapped length | 2.7 Gb | 2.7 Gb |
| Number of chromosomes | 21 | 21 |
| Number of scaffolds | 41 | 101 |
| Scaffold N50 | 144.4 Mb | 106.1 Mb |
| Scaffold L50 | 9 | 11 |
| Number of contigs | 8,743 | 305 |
| Contig N50 | 40.8 Mb | 59.5 Mb |
| Contig L50 | 23 | 15 |
| GC percent | 41 | 41.5 |

### Annotation

The preliminary annotation file along with the annotation comparison between B10.RIII and GRCm39 is provided as the supplementary file.

### Conclusion

In this study, we generated the first, high-quality chromosome level genome sequence of B10.RIII, using a combination of long read, short read and optical mapping techniques. The B10.RIII genome sequence scored higher than the reference mouse genome both in assembly completeness and assembly quality. The B10.RIII genome annotation based on the reference genome identified about 98% of the known reference genes. We believe the availability of B10.RIII genome sequence will help advance the understanding of autoimmune diseases, giving the researchers a better idea on its unique genetic makeup. Comparative studies of B10.RIII with other mouse models could also significantly enhance the power of these models towards their clinical applications.

## Data availability

All of the raw data have been deposited in public repositories and are freely available from NCBI using the following accessions.

BioSample: SAMN26583198 (https://www.ncbi.nlm.nih.gov/biosample/SAMN26583198) BioProject: PRJNA814472 (https://www.ncbi.nlm.nih.gov/bioproject/PRJNA814472)

SRA Illumina: SRR18305688

SRA pacbio: SRR18290958, SRR18500559, SRR18500560, SRR18500558 SRA bionano: SUPPF_0000004450

The B10.RIII genome assembly is available from the NCBI using the following accession; Genome: GCA_030265425.1

(https://www.ncbi.nlm.nih.gov/datasets/genome/GCA_030265425.1/)

All the code/commands used in this study are freely available to the public, through the GitHub repository at; https://github.com/NIH-NEI/b10riii-genome-assembly

## Abbreviations

AID: Autoimmune Disease
RA: Rheumatoid Arthritis
SLE: Systemic Lupus Erythematosus
AS: Ankylosing Spondylitis
EAE: Experimental Autoimmune Encephalomyelitis
EAU: Experimental Autoimmune Uveitis
EAN: Experimental Autoimmune Neuritis
EAT: Experimental Autoimmune Thyroiditis
ECIA: Experimental Collagen-Induced Arthritis
WGS: Whole Genome Sequencing
gDNA: Genomic DNA

## Additional files

1. genomescope_summary.txt – Genomescope output file.
2. jellyhisto.txt – Jellyfish output file.
3. kmergenie_histograms_report.html – KmerGenie output file.
4. reference-b10-annotation-comparison.xlsx – Annotation comparison
5. b10.riii.gff3 – B10.RIII reference based annotation file

## Disclosure of use of AI-assisted tools including generative AI

No content generated by any AI-assisted tools are used in this manuscript.

## Ethics

This study was approved by the NIH Animal Protocol Study (APS) Number NEI-688.

## Acknowledgements

This work utilized the computational resources of the NIH HPC Biowulf cluster. The library preparation and sequencing were carried out at the NCI CCR Sequencing Core Facility.

## Authors contributions

VN, YJ, RH and RC conceived and designed the study. YJ and RH performed the experiments. VN analyzed the data. VN, YJ and GS drafted the manuscript. VN, YJ, GS, RH and RC critically reviewed the manuscript.

## Software Versions Used

bamtools 2.5.2

bwa 0.7.17

bbtools 38.87

fastqc 0.11.9

GenomeScope 1.0.0

gcpp 2.0.2

HiCanu 2.2

Hifiasm 0.16.1

HiFiAdapterFilt 2.0.0

jellyfish 2.3.0

KmerGenie 1.7048

pbmm2 1.9.0

pbipa 1.5.0

pilon 1.24

samtools 1.17

MaSuRCA 4.1.0

QUAST 5.2.0

gffcompare 0.12.6

bionano HybridScaffold 1.0

[TAB]

## Reference

1. Jacobson, D.L., et al., Epidemiology and estimated population burden of selected autoimmune diseases in the United States. Clin Immunol Immunopathol, 1997. 84(3): p. 223–43.

2. Fugger, L., L.T. Jensen, and J. Rossjohn, Challenges, Progress, and Prospects of Developing Therapies to Treat Autoimmune Diseases. Cell, 2020. 181(1): p. 63–80.

3. Bieber, K., et al., Autoimmune pre-disease. Autoimmun Rev, 2023. 22(2): p. 103236.

4. Lee, D.S.W., O.L. Rojas, and J.L. Gommerman, B cell depletion therapies in autoimmune disease: advances and mechanistic insights. Nat Rev Drug Discov, 2021. 20(3): p. 179–199.

5. Guenet, J.L., The mouse genome. Genome Res, 2005. 15(12): p. 1729–40.

6. Billiau, A. and P. Matthys, Modes of action of Freund’s adjuvants in experimental models of autoimmune diseases. J Leukoc Biol, 2001. 70(6): p. 849–60.

7. Krishnamoorthy, G. and H. Wekerle, EAE: an immunologist’s magic eye. Eur J Immunol, 2009. 39(8): p. 2031–5.

8. Caspi, R.R., et al., Mouse models of experimental autoimmune uveitis. Ophthalmic Res, 2008. 40(3-4): p. 169–74.

9. Chan, C.C., et al., Pathology of experimental autoimmune uveoretinitis in mice. J Autoimmun, 1990. 3(3): p. 247–55.

10. Maurer, M. and R. Gold, Animal models of immune-mediated neuropathies. Curr Opin Neurol, 2002. 15(5): p. 617–22.

11. Kong, Y.M., Experimental autoimmune thyroiditis in the mouse. Curr Protoc Immunol, 2007. Chapter 15: p. 15 7 1–15 7 21.

12. Cho, Y.G., et al., Type II collagen autoimmunity in a mouse model of human rheumatoid arthritis. Autoimmun Rev, 2007. 7(1): p. 65–70.

13. Klaska, I.P. and J.V. Forrester, Mouse models of autoimmune uveitis. Curr Pharm Des, 2015. 21(18): p. 2453–67.

14. Jansson, L., et al., Chronic experimental autoimmune encephalomyelitis induced by the 89-101 myelin basic protein peptide in B10RIII (H-2r) mice. Eur J Immunol, 1991. 21(3): p. 693–9.

15. Jirholt, J., et al., Genetic linkage analysis of collagen-induced arthritis in the mouse. Eur J Immunol, 1998. 28(10): p. 3321–8.

16. Caspi, R.R., et al., Genetic control of susceptibility to experimental autoimmune uveoretinitis in the mouse model. Concomitant regulation by MHC and non-MHC genes. J Immunol, 1992. 148(8): p. 2384–9.

17. Sarsani, V.K., et al., The Genome of C57BL/6J “Eve”, the Mother of the Laboratory Mouse Genome Reference Strain. G3 (Bethesda), 2019. 9(6): p. 1795–1805.

18. Church, D.M., et al., Lineage-specific biology revealed by a finished genome assembly of the mouse. PLoS Biol, 2009. 7(5): p. e1000112.

19. Lilue, J., et al., Sixteen diverse laboratory mouse reference genomes define strain-specific haplotypes and novel functional loci. Nat Genet, 2018. 50(11): p. 1574–1583.

20. Chikhi, R. and P. Medvedev, Informed and automated k-mer size selection for genome assembly. Bioinformatics, 2014. 30(1): p. 31–7.

21. Marcais, G. and C. Kingsford, A fast, lock-free approach for efficient parallel counting of occurrences of k-mers. Bioinformatics, 2011. 27(6): p. 764–70.

22. Vurture, G.W., et al., GenomeScope: fast reference-free genome profiling from short reads. Bioinformatics, 2017. 33(14): p. 2202–2204.

23. Andrews, S., FastQC : A quality control tool for high throughput sequence data. 2023.

24. Bushnell, B., BBDUK : Decontamination Using Kmers. 2021.

25. Sim, S.B., et al., HiFiAdapterFilt, a memory efficient read processing pipeline, prevents occurrence of adapter sequence in PacBio HiFi reads and their negative impacts on genome assembly. BMC Genomics, 2022. 23(1): p. 157.

26. Nurk, S., et al., HiCanu: accurate assembly of segmental duplications, satellites, and allelic variants from high-fidelity long reads. Genome Res, 2020. 30(9): p. 1291–1305.

27. Biosciences, P., IPA: Improved Phased Assembly. 2022.

28. Cheng, H., et al., Haplotype-resolved de novo assembly using phased assembly graphs with hifiasm. Nat Methods, 2021. 18(2): p. 170–175.

29. Biosciences, P. pbmm2 : A minimap2 SMRT wrapper for PacBio data. 2023; Available from: https://github.com/PacificBiosciences/pbmm2.

30. Li, H., et al., The Sequence Alignment/Map format and SAMtools. Bioinformatics, 2009. 25(16): p. 2078–9.

31. Derek Barnett, E.G., Gabor Marth, Michael Stromberg. BamTools. 2009; Available from: https://github.com/pezmaster31/bamtools.

32. Biosciences, P. GCpp: Generate Highly Accurate Reference Contigs. 2023; Available from: https://github.com/PacificBiosciences/gcpp.

33. Walker, B.J., et al., Pilon: an integrated tool for comprehensive microbial variant detection and genome assembly improvement. PLoS One, 2014. 9(11): p. e112963.

34. Bionano, HybridScaffold. 2022.

35. Zimin, A.V., et al., The MaSuRCA genome assembler. Bioinformatics, 2013. 29(21): p. 2669–77.

36. Alonge, M., et al., Automated assembly scaffolding using RagTag elevates a new tomato system for high-throughput genome editing. Genome Biol, 2022. 23(1): p. 258.

37. Shumate, A. and S. Salzberg, LiftoffTools: a toolkit for comparing gene annotations mapped between genome assemblies. F1000Res, 2022. 11: p. 1230.

38. J, D., AGAT: Another Gff Analysis Toolkit to handle annotations in any GTF/GFF format. 2024: Zenodo.

39. Gurevich, A., et al., QUAST: quality assessment tool for genome assemblies. Bioinformatics, 2013. 29(8): p. 1072–5.

