## Supplementary material for "Chromosome-scale genome assembly of B10.RIII, an autoimmune susceptible mouse strain": kmergenie_histograms_report.html

### KmerGenie report

#### Predicted best k: 117

###### Predicted assembly size: 2432633107 bp

The above plot should be roughly concave and have a clear global maximum. If not, the predicted best k is likely to be inaccurate.  
Click here for more details.

This box will guide you towards analyzing the main plot produced by Kmergenie (# genomic k-mers vs k).

The y axis is a number of k-mers, but can also be interpreted as the the estimated genome size (in bp) when repeats are collapsed, i.e. the estimated size of an assembly of this dataset.
Note that, according to the Kmergenie article, this plot should be roughly concave.

Thus, in the ideal case, the above plot should be a smooth curve with a clear global maximum, i.e.:

  

However, in some cases, the plot may be more similar to these, with multiple local maximas or a plateau:

  

These cases reflect that the statistical model in Kmergenie does not always correctly fit the input data for some values of k. Thus, the best k value predicted by Kmergenie may be suboptimal. In your subsequent analyses (e.g. de novo assembly), we recommend that you also try a larger k than the one predicted by Kmergenie, following the indications in green in the above plots. Essentially, when the number of estimated genomic k-mers remains high during a large range of k's, the largest k value in this range is likely to be a better choice.

In other cases, the number of genomic k-mers never drops with high k values. This is an indication of a highly covered dataset:

In this case, it is recommended to restart Kmergenie with a higher maximal k value (higher than the default value of 120). An unusually high k value (>100) may produce good results.

Hide this box

##### Sampled histogram and fit for each value of k

Colors of the fits: red is the fit of the complete statistical model of the histogram (erronous k-mers + genomic k-mers). When using the diploid model, blue are only the heterozygous k-mers, green are only the homozygous k-mers.


Generated by KmerGenie
